# Elucidating effects of single and multiple resistance mechanisms on bacterial response to meropenem by quantitative and systems pharmacology modeling and population genomics

**DOI:** 10.1101/2024.02.17.579784

**Authors:** Dominika T. Fuhs, Sara Cortés-Lara, Jessica R. Tait, Kate E. Rogers, Carla López-Causapé, Wee Leng Lee, David M. Shackleford, Roger L. Nation, Antonio Oliver, Cornelia B. Landersdorfer

**Affiliations:** Drug Delivery, Disposition and Dynamics, Monash Institute of Pharmaceutical Sciences, Monash University, Parkville, Victoria, Australia; Servicio de Microbiología, Hospital Universitario Son Espases, Instituto de Investigación Sanitaria Illes Balears (IdISBa), Palma de Mallorca, Spain; CIBER Enfermedades Infecciosas (CIBERINFEC), Madrid, Spain; Centre for Drug Candidate Optimisation, Monash Institute of Pharmaceutical Sciences, Monash University, Parkville, Victoria, Australia

## Abstract

Meropenem is commonly used against *Pseudomonas aeruginosa*. Traditionally, the time unbound antibiotic concentration exceeds the MIC (*f*T_>MIC_) is used to select carbapenem regimens. We aimed to: characterize the effects of different baseline resistance mechanisms on bacterial killing and resistance emergence; evaluate whether *f*T_>MIC_ can predict these effects; and, develop a novel quantitative and systems pharmacology (QSP) model to describe effects of baseline resistance mechanisms on the time-course of bacterial response.

Seven isogenic *P. aeruginosa* strains with a range of resistance mechanisms and MICs were used in 10-day hollow-fiber infection model studies. Meropenem pharmacokinetic profiles were simulated for various regimens (t_1/2,meropenem_=1.5h). All viable counts on drug-free, 3×MIC and 5×MIC meropenem-containing agar across all strains, five regimens and control (n=90 profiles) were simultaneously subjected to QSP modeling. Whole genome sequencing was completed for total population samples and emergent resistant colonies at 239h.

Regimens achieving ≥98%*f*T_>1xMIC_ suppressed resistance emergence of the *mexR* knockout strain. Even 100%*f*T_>5xMIC_ failed to achieve this against the strain with OprD loss and the *ampD* and *mexR* double-knockout strain. Baseline resistance mechanisms affected bacterial outcomes, even for strains with the same MIC. Genomic analysis revealed that pre-existing resistant subpopulations drove resistance emergence. During meropenem exposure, mutations in *mexR* were selected in strains with baseline *oprD* mutations, and *vice versa*, confirming these as major mechanisms of resistance emergence. Secondary mutations occurred in *lysS* or *argS*, coding for lysyl and arginyl tRNA synthetases, respectively. The QSP model well characterized all bacterial outcomes of the seven strains simultaneously, which *f*T_>MIC_ could not.

## Introduction

Carbapenem-resistant *Pseudomonas aeruginosa* is prevalent in intensive care units (ICUs), causing high mortality and morbidity (1, 2). The World Health Organization identified carbapenem-resistant *P. aeruginosa* as a top three critical pathogen (3). In 2019, *P. aeruginosa* was among the six leading pathogens causing deaths associated with antimicrobial resistance worldwide (4). *P. aeruginosa* possesses an exceptional ability for resistance emergence during antibiotic treatment (5, 6).

Antibiotic development cannot keep up with the rise in antimicrobial resistance and resistance emerges rapidly even against new compounds (7, 8). Therefore, there is a pressing need for new approaches to describe and predict resistance emergence during antibiotic exposure and optimise antibiotic dosing regimens (9). Traditionally, pharmacokinetic/pharmacodynamic (PK/PD) indices have linked antibiotic exposure with bacterial response and informed dosing (10, 11). For example, for carbapenems the relevant PK/PD index is the percentage of time during which unbound antibiotic concentrations exceed the minimum inhibitory concentration (MIC) over 24h (%*f*T_>MIC_) (12). However, limitations of these indices are increasingly recognized, including that they do not capture the full time-courses of antibiotic PK and bacterial response, and do not appropriately account for resistance emergence. Other limitations relate to their use of MIC as a PD measure of drug potency (9, 13). In contrast, mechanism-based PK/PD and Quantitative and Systems Pharmacology (QSP) models describe and predict the full time-courses of bacterial growth, killing and resistance emergence (14, 15).

Numerous genes and mutations are involved in *P. aeruginosa* resistance (16, 17). Carbapenem resistance may be attributed to OprD porin channel inactivation, hyperproduction of efflux pumps such as MexAB-OprM, production of antibiotic-deactivating enzymes, or combinations thereof (18). Currently, there is a lack of knowledge on the impact of various *P. aeruginosa* baseline resistance mechanisms, *i.e.* that are present pre-treatment, on bacterial killing and resistance emergence during treatment. Since meropenem is commonly used in ICUs against *P. aeruginosa*, and carbapenem-resistant strains remain a major health problem, we have used meropenem as a probe to explore this issue (1, 19, 20). Published meropenem PK/PD models for *P. aeruginosa* did not directly incorporate effects of baseline bacterial characteristics on resistance emergence (21-33). To the best of our knowledge no studies have employed an integrated experimental and modeling approach to systematically investigate meropenem regimens in the context of different resistance mechanisms using a panel of isogenic strains.

Therefore, the objectives of this study were to (i) characterize the effect of different meropenem regimens on bacterial killing and resistance emergence in seven isogenic *P. aeruginosa* strains with one or multiple baseline resistance mechanisms, (ii) identify genetic changes in bacteria relating to resistance emergence during meropenem exposure, (iii) assess whether the observed bacterial responses of the strains are consistent with the relevant traditional PK/PD index (%*f*T_>MIC_), and (iv) develop a QSP model that quantitatively describes the effect of baseline resistance mechanisms on bacterial killing and resistance emergence across strains.

## Materials and methods

### Antibiotic, media, strains and susceptibility testing

Stock solutions of meropenem (Fresenius Kabi, New South Wales, Australia) were prepared in Milli-Q water, sterilized using a 0.22µm PVDF filter and stored at -80°C. All studies used cation-adjusted Mueller Hinton broth (CAMHB; BD, Sparks, MD, USA) containing 20-25mg/L Ca^2+^ and 10-12.5mg/L Mg^2+^. Total viable counting was performed on cation-adjusted Mueller Hinton agar (CAMHA; BD, Sparks, MD, USA) containing 20-25mg/L Ca^2+^ and 10-12.5mg/L Mg^2+^. Meropenem-containing plates were prepared by supplementing CAMHA with appropriate volumes of drug stock solution. The *P. aeruginosa* wild-type reference strain PAO1 and six isogenic strains constructed from PAO1 (PAΔAD, PAOD1, PAΔmexR, PAΔDΔmexR, PAOD1ΔmexR and PAOD1ΔD) were studied and are described in **Table 1**. The only differences between PAO1 and its isogenic strains were the introduced resistance mechanisms. This allowed for direct investigation of the impact of these resistance mechanisms (by themselves or in combinations of two) on bacterial response. The meropenem MIC prior to antibiotic exposure (**Table 1**) and baseline log_10_ mutant frequency (MF) were determined for all strains in biological replicates on different days (**Table S1**), as described in the **Supplementary materials**.

**Table 1:** Isogenic strains of *P. aeruginosa* and their baseline MICs.

| Strain | Description | MIC<sub>MEM</sub><br>(mg/L)* | Susceptibility<br>to MEM<sup>#</sup> | Ref. |
| --- | --- | --- | --- | --- |
| PAO1 | Wild-type reference strain | 1 | S | - |
| PAΔAD | *ampD* knockout resulting in AmpC overexpression | 2 | S | (62) |
| PAΔmexR | *mexR* knockout resulting in overexpression of MexAB-OprM efflux | 4 | I | (59) |
| PAOD1 | Spontaneous *oprD* mutant resulting in the loss of OprD porin channels | 4 | I | (63) |
| PAΔDΔmexR | *ampD* and *mexR* knockout | 4 | I | (59) |
| PAOD1ΔmexR | Spontaneous *oprD* mutant and *mexR* knockout | 16 | R | (59) |
| PAOD1ΔD | Spontaneous *oprD* mutant and *ampD* knockout | 16 | R | (63) |

Δ: deletion, MEM: meropenem, \*determined by agar dilution (64) <sup>#</sup>according to CLSI clinical breakpoints, S: susceptible, I: intermediate, R: resistant. All isogenic strains were constructed from PAO1 by the group of Antonio Oliver, Instituto de Investigación Sanitaria Illes Balears.

### Dynamic hollow-fiber infection model (HFIM) studies

Free (non-protein bound) meropenem concentration-time profiles were simulated *in silico* using Berkeley Madonna (v8.3.18), based on published population PK studies in critically ill patients with normal renal function (CL_CR_ 120mL/min, meropenem t_1/2_ 1.5h) (34, 35), using clinically-relevant regimens (and associated modes of administration) as summarized in **Table 2**. Regimens were 1g and 2g as 3-h infusions given 8-hourly (Q8h), and 3g/day, 6g/day and 12g/day as continuous infusions (CI) (19, 36-38). Untreated controls were included for each strain. **Table 2** summarizes all regimens and biological replicates. Dosing regimens were selected based on clinical relevance and included different modes of administration. The higher than approved 12g/day CI was included to reflect meropenem doses and concentrations related to high-dose regimens and to further explore the concentration-effect relationship for the meropenem-resistant strains (39, 40).

**Table 2:** Meropenem (MEM) regimens (and mode of administration) simulated in the hollow-fiber infection model against seven isogenic *P. aeruginosa* strains.

| MEM regimens | Isogenic strains (number of biological replicates) |
| --- | --- |
| 1 g, Q8h | PAO1 (1), PAΔAD (1), PAΔmexR (1), PAOD1 (1), PAΔDΔmexR (1) |
| 2 g, Q8h | PAΔmexR (1), PAOD1 (1), PAΔDΔmexR (1) |
| 3 g/day, CI | PAO1 (1), PAΔAD (1), PAΔmexR (1), PAOD1 (1), PAΔDΔmexR (1) |
| 6 g/day, CI | PAΔmexR (1), PAOD1 (2), PAΔDΔmexR (3), PAOD1ΔmexR (1), PAOD1ΔD (2) |
| 12 g/day, CI | PAΔmexR (1), PAOD1 (1), PAΔDΔmexR (1), PAOD1ΔmexR (1), PAOD1ΔD (2) |

Q8h, 3-h infusion administered 8-hourly; CI, continuous infusion.
Since the standard 1 g Q8h and 3 g daily CI meropenem dosing regimens resulted in bacterial killing and suppression of emergence of resistance for PAO1 and PAΔAD strains, higher meropenem dosing regimens were not tested against these strains.

The HFIM studies utilized cellulosic cartridges (C3008 B; FiberCell Systems Inc., Frederick, MD, USA) in a humidified incubator at 37°C for 10 days, as previously described (27, 41, 42). In brief, for each strain, a culture of 3 colonies was grown overnight in CAMHB at 37°C, and the optical density measured spectrophotometrically. The bacterial suspension was then diluted to achieve a targeted initial inoculum of 10^7^CFU/mL and 17mL were injected into the HFIM cartridge. The Q8h regimens were delivered via syringe drivers into the central reservoir of the HFIM. Each CI was initiated with an appropriate loading dose at 0h as a bolus into the central reservoir to achieve the targeted steady-state concentration. The CI concentrations were maintained by adding meropenem into the fresh media bottles flowing to the central reservoir. The media bottles were stored in the fridge and replaced every 48h. Media samples for PK validation (by LC-MS analysis per **Supplementary materials**; 1mL in duplicates) were collected in cryovials throughout the experiment from the outflow of the central reservoir and immediately stored at -80°C.

Bacterial samples (1.5mL) were collected at 0, 3.5, 7, 12.5, 23, 27.5, 31, 47, 51.5, 55, 71, 95, 119, 143, 167, 191, 215, and 239h. Samples were twice centrifuged (4,000×*g*, 5min), resuspended in saline to reduce antibiotic carryover and manually plated on CAMHA containing no drug (100µL plated, all time points) and meropenem-containing CAMHA plates at 3× and 5×MIC (200µL plated, 0, 23, 47, 71, 119, 167, 215 and 239h). Bacterial samples plated on drug-free plates were incubated for 24h at 37°C with a limit of counting of 1.0 log_10_CFU/mL. Bacterial samples plated on drug-containing agar were incubated for 48-72h at 37°C with a limit of counting of 0.7 log_10_CFU/mL. Bacterial colonies that grew on meropenem-containing plates were selected and stored at -80°C for post-antibiotic exposure MIC testing and genomic analysis. Population samples for genome sequencing were collected from the cartridge at 239h, centrifuged (4000×*g*, 5min), the supernatant removed and pellets stored at -80°C.

### Whole genome sequencing (WGS) and analysis of emergent meropenem resistance mechanisms

Total population samples and saved colonies that grew on 5xMIC meropenem-containing CAMHA from each isogenic strain and dosing regimen (including the untreated arms) obtained at 239h, were fully sequenced. The population samples and colonies from PAO1 were also sequenced. Details are described in the **Supplementary materials**.

### Quantitative and systems pharmacology modeling

A QSP model was developed to account for bacterial growth, killing and resistance emergence for all five regimens and seven strains simultaneously, importantly without estimating strain-specific drug effect parameters. The model was informed by the total and less-susceptible bacterial counts, baseline resistance mechanisms and genome sequencing. It explained the bacterial outcomes by incorporating effects of different resistance mechanisms on the estimated meropenem concentration at the site of action (periplasmic space) which drives bacterial response. See the **Supplementary materials** for details, including a schematic diagram (**Figure S1**).

## Results

### MICs and baseline mutant frequencies

The studied strains encompassed a wide range of meropenem susceptibilities (**Table 1**). The baseline log_10_MFs indicated presence of a small proportion of pre-existing less-susceptible bacterial subpopulations in the overall population of each strain (**Table S1**).

### Dynamic hollow-fiber infection model studies

The observed meropenem concentrations for CI regimens were on average within 2% of targeted concentrations. The targeted meropenem concentrations for the Q8h regimens were also well replicated (**Figure S2**). All strains grew well in the control arms, reaching ∼10^10^CFU/mL by 24h which were maintained throughout the experiments. Changes in total and less-susceptible bacterial populations growing on 3× and 5×MIC meropenem-containing CAMHA are shown in **Figures 1,2,3**, respectively. The less-susceptible populations in the untreated controls plateaued at ∼3-4 log_10_CFU/mL by ∼24h, except for PAO1 where the plateau occurred from ∼48h (**Figures 2,3**).

**Figure 1:**
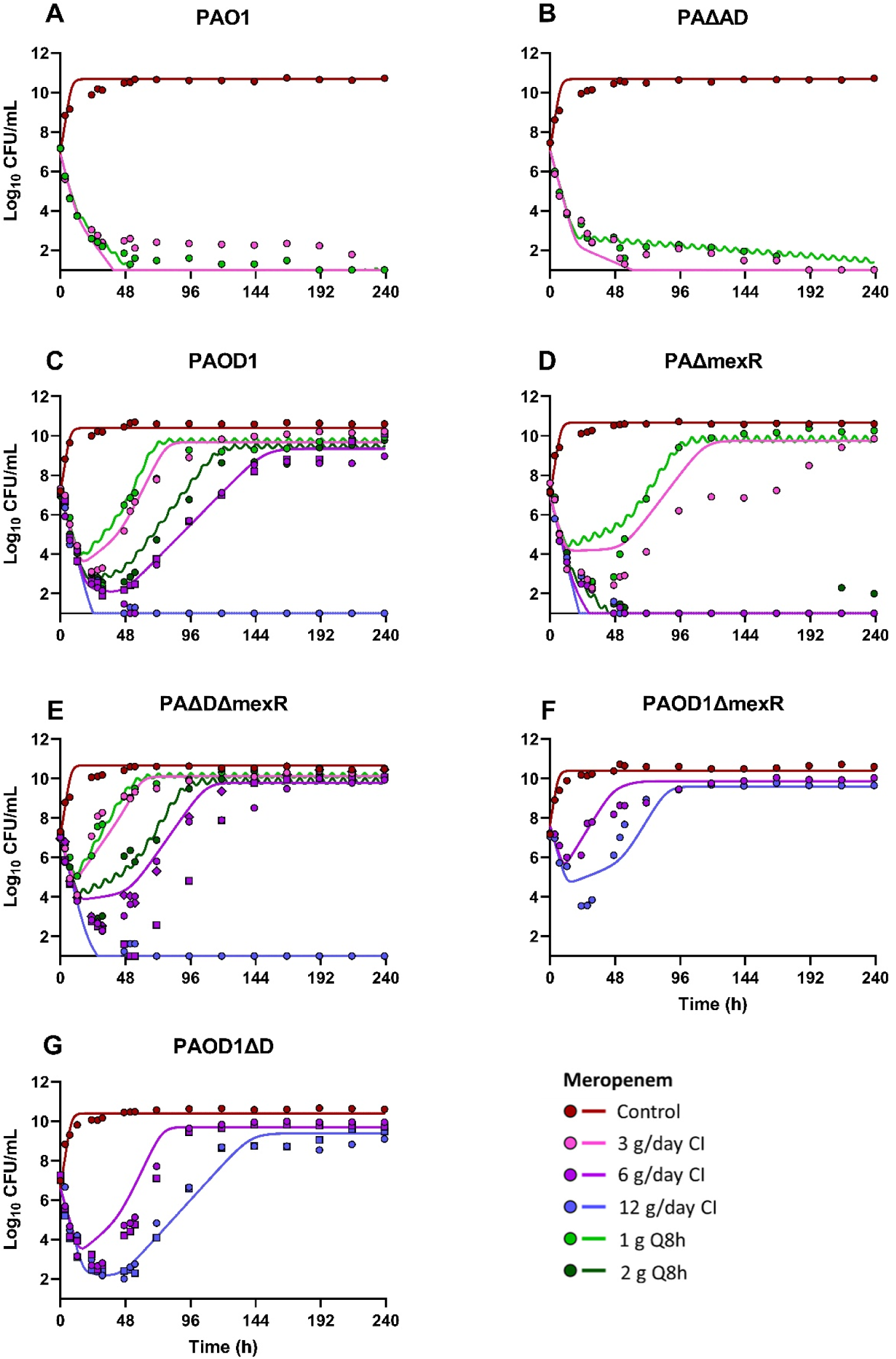
Total bacterial population (observed viable counts in symbols, and population predicted profiles of the mechanism-based model in lines in corresponding colours) vs time. The y axis starts at the limit of counting = 1 log_10_CFU/mL. n=1-3, biological replicates are shown as different symbols.

**Figure 2:**
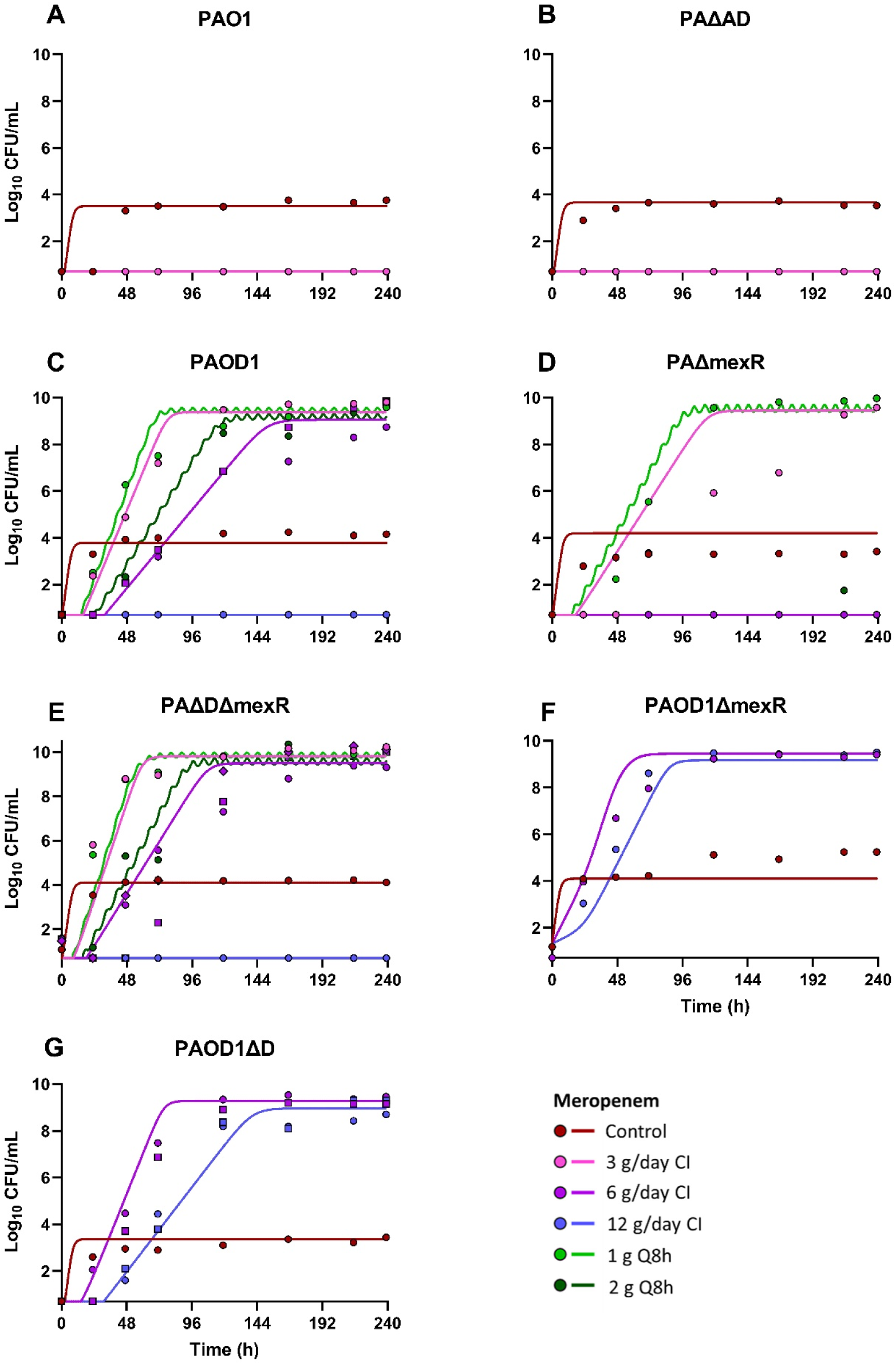
Less-susceptible bacterial population growing on 3× MIC MEM-containing drug plates (observed viable counts in symbols, and population predicted profiles of the mechanism-based model in lines in corresponding colours) vs time. The y axis starts at the limit of counting = 0.7 log_10_CFU/mL. n=1-3, biological replicates are shown as different symbols.

**Figure 3:**
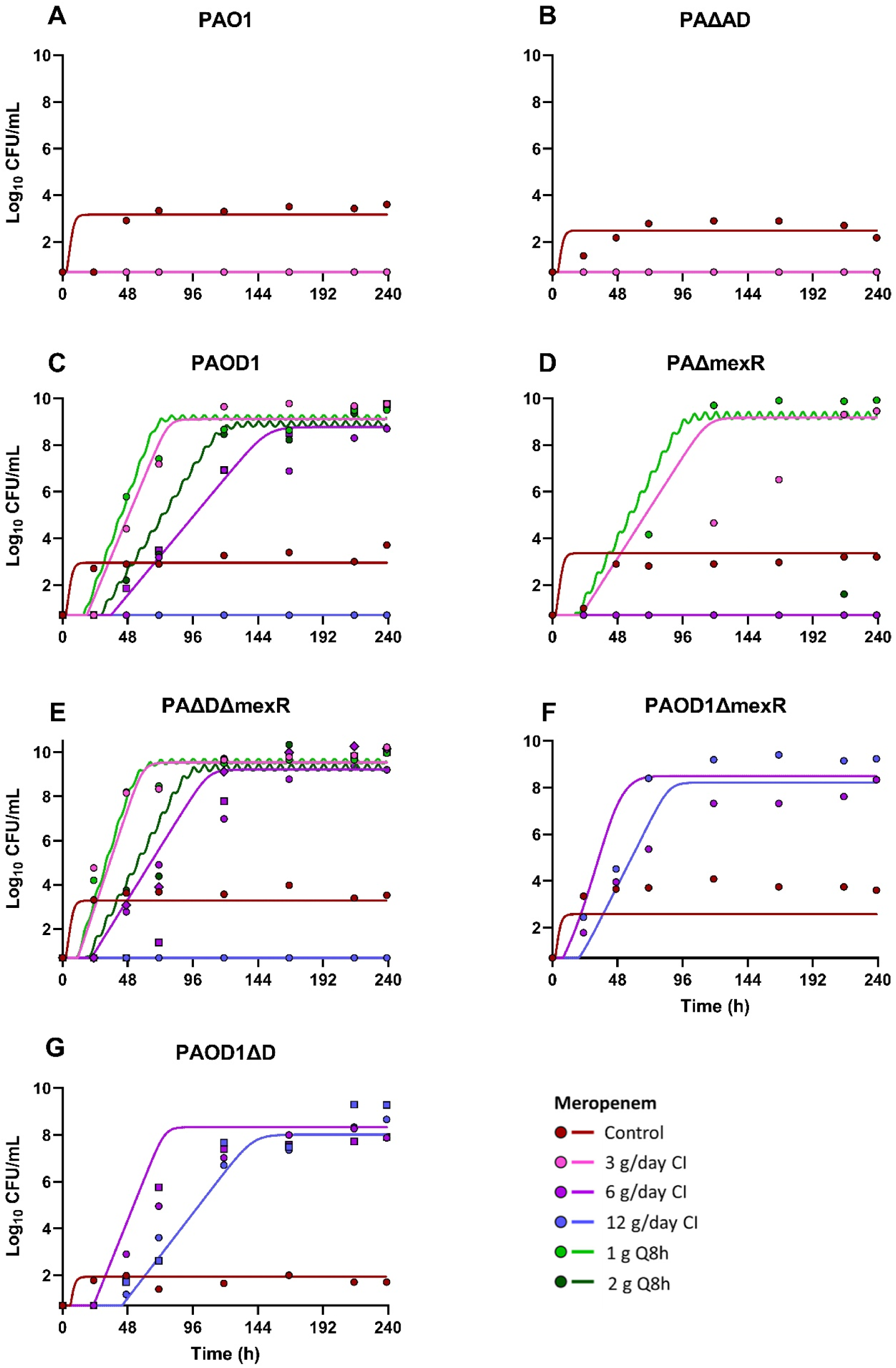
Less-susceptible bacterial population growing on 5× MIC MEM-containing drug plates (observed viable counts in symbols, and population predicted profiles of the mechanism-based model in lines in corresponding colours) vs time. The y axis starts at the limit of counting = 0.7 log_10_CFU/mL. n=1-3, biological replicates are shown as different symbols.

#### PAO1 and PAΔAD

Exposure to 3g/day CI and 1g Q8h regimens (≥98% *f*T_>MIC_) resulted in ≥5 log_10_CFU/mL killing and suppressed resistance emergence (**Figure 1A,B**). Less-susceptible bacteria were not detected throughout the experiments for both strains (**Figures 2A,B,3A,B**). No effect of the *ampD* mutation was observed when comparing PAΔAD to PAO1, despite slightly different baseline MICs.

#### PAΔmexR, PAOD1 and PAΔDΔmexR

The lowest CI and Q8h regimens resulted in bacterial regrowth with resistance emergence for all three strains (**Figure 1C,D,E**), although the CI achieved 100% *f*T_>2xMIC_. Regrowth of the PAΔDΔmexR double mutant occurred substantially faster (from ∼23h) compared to the other two strains. The 6g/day CI (100% *f*T_>5xMIC_) and 2g Q8h (98% *f*T_>MIC_) regimens produced ≥5 log_10_CFU/mL killing and supressed the regrowth of less-susceptible bacteria only against PAΔmexR. Despite having the same MIC as PAΔmexR, the PAOD1 and PAΔDΔmexR strains displayed extensive regrowth associated with resistance emergence (**Figures 2C,E,3C,E**). Addition of the *ampD* mutation to the *mexR* mutation in PAΔDΔmexR substantially affected bacterial response, as compared to PAΔmexR. In PAΔDΔmexR, the two resistance mechanisms acted synergistically in decreasing the meropenem effect. Only the highest (12g/day) CI regimen, that achieved 100% *f*T_>10×MIC_, produced bacterial eradication and suppressed resistance emergence of all three strains.

#### PAOD1ΔmexR and PAOD1ΔD

For both strains, the 6 and 12g/day CI regimens, resulted in extensive regrowth and resistance emergence (**Figures 1,2,3 F,G**). The 6 g/day CI (100% *f*T_>MIC_) achieved only ∼1 log_10_CFU/mL killing of PAOD1ΔmexR followed by almost instantaneous regrowth with resistance emergence (**Figures 1,2,3F**). The same regimen produced ∼5 log_10_CFU/mL initial bacterial killing of PAOD1ΔD (**Figures 1,2,3G**) before regrowth of less-susceptible bacteria, although both strains had the same meropenem MIC (16mg/L). Even 12g/day CI (100% *f*T_>2xMIC_) was unable to suppress regrowth and resistance emergence of either strain.

#### All strains

All treatments resulting in regrowth amplified resistance with bacterial counts greater than the baseline resistance shown in control populations (**Figures 2,3**). Overall, 3-h infusions Q8h performed similarly to CI at a given daily dose, with trends of a slightly better performance of the 3g CI against PAΔmexR, and the 6g CI against PAOD1 and PAΔDΔmexR.

### Genome sequencing and analysis of emergent meropenem resistance mechanisms

The mutations detected at 239h in total population samples and individual colonies that grew on meropenem-containing CAMHA are presented in **Table 3**. All analysed colonies from meropenem-containing CAMHA retained resistance (supported by significantly increased meropenem MICs, **Table 3**) after being cultured in drug-free media. Following the HFIM studies, several of the studied isogenic strains exhibited additional mutations in genes known to be responsible for meropenem resistance. In general, mutations in *mexAB* regulators (resulting in MexAB-OprM efflux pump overexpression) were detected in isogenic strains with baseline *oprD* mutation. Mutations in the *oprD* gene (resulting in OprD porin inactivation) were detected in isogenic strains with baseline *mexR* mutations. Secondary mutations occurred mostly in *lysS* or *argS*, coding for lysyl and arginyl tRNA synthetases, respectively. Other genes found to be mutated in more than one experiment included *alaS* (alanyl-tRNA synthetase), *phoQ* (two-component sensor PhoQ), *spoT* (guanosine-3’,5’-bis(diphosphate) 3’-pyrophosphohydrolase) and *aroB* (3-dehydroquinate synthase). Finally, a large genomic deletion of *galU* region was detected in a single mutant colony from a control PAΔmexR experiment and a PBP3 (F533L) mutation in a single mutant colony from a 1g Q8h meropenem treatment of PAΔDΔmexR.

**Table 3:**
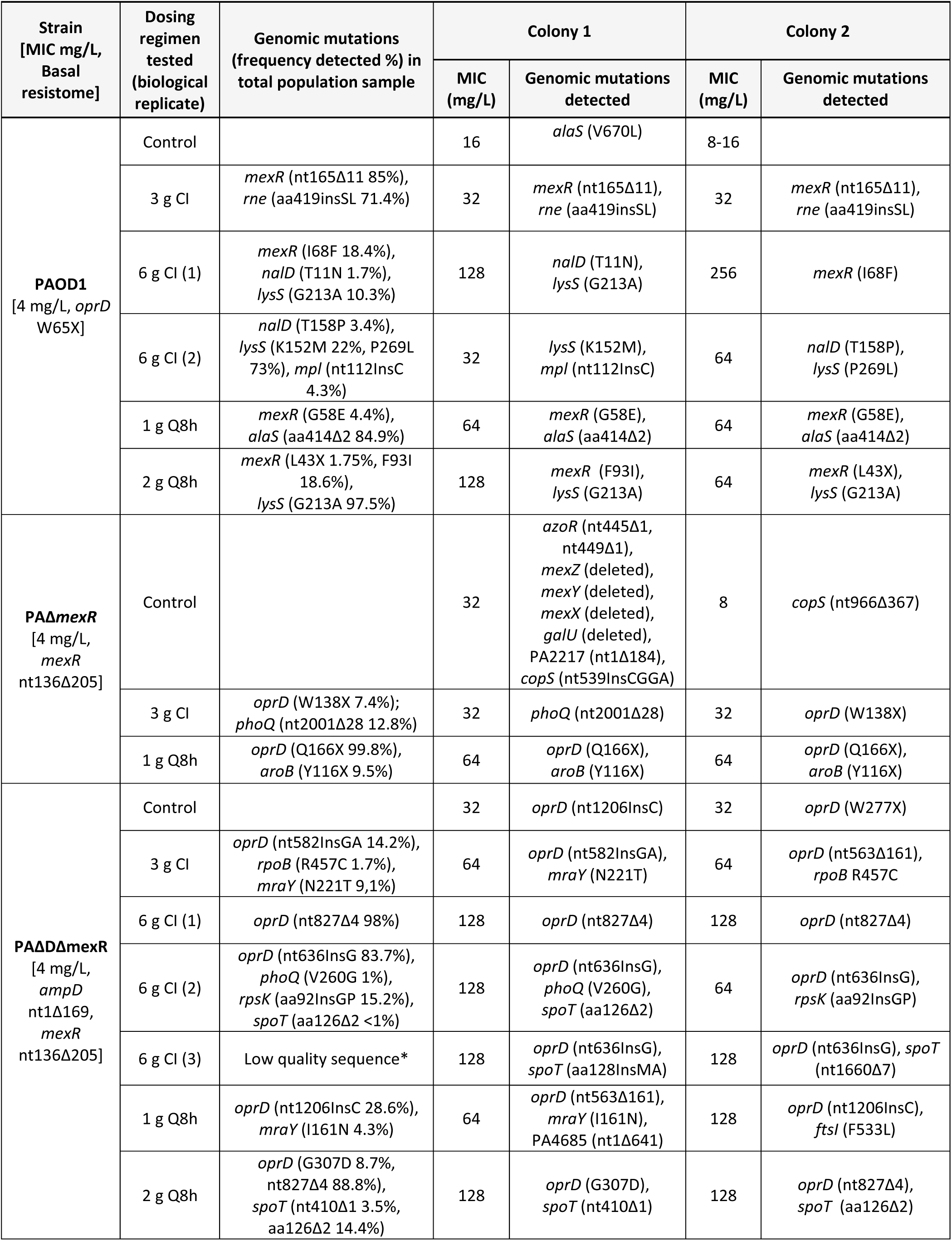
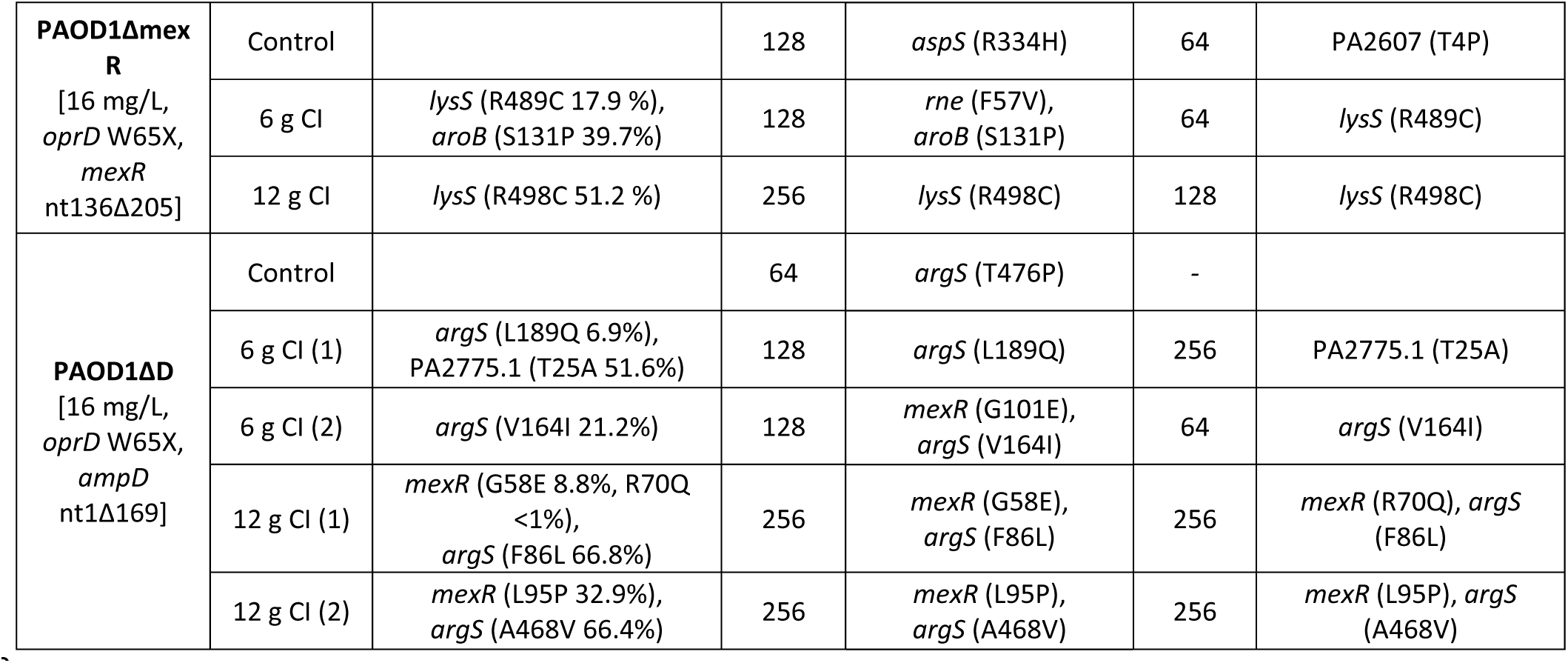
Mutations detected in total population sample and saved colonies of isogenic strains of *P. aeruginosa* after the completion of HFIM (239 h).

### QSP modeling

In this model, amplification of small pre-existing resistant bacterial subpopulations (CFU_R_) that had additional resistance mutations compared to the baseline resistome of the respective strain, was responsible for driving the observed bacterial regrowth and resistance emergence, supported by the genomic analysis. The model yielded unbiased and sufficiently precise fits for the total bacterial counts for all strains (**Figure 1**), as well as the growth and killing of less-susceptible populations (**Figures 2,3**), demonstrated by the population fitted curves that do not include any random variability. The coefficient of correlation was 0.98 for observed *versus* individual-fitted counts and 0.94 for observed *versus* population-fitted counts (**Figure S3A,B**). The observed *versus* individual-fitted and observed *versus* population-fitted plots also showed highly sufficient performance for the less-susceptible populations (**Figure S3C,D**). The data from all tested regimens and controls across all seven strains were described simultaneously by the same model and largely same parameter estimates (**Table 4**). Distinct parameter estimates were required only to describe the observed differences in initial inocula, and the proportions of pre-existing bacterial subpopulations between strains (**Table 5**). The differences in bacterial killing and resistance emergence among strains were characterized mainly by the effects of a given resistance mechanism/s present in the strains. All standard errors (SE) of parameter estimates were <16%, indicating excellent precision. The model very well described up to five different treatment and control arms for each of seven isogenic strains and their biological replicates (including total and less-susceptible subpopulations), *i.e.* in total 90 curves modeled simultaneously. The PAO1 strain, challenged with 3g/day CI, had a low concentration of total bacteria (∼2 log_10_CFU/mL) remaining until 215h (**Figure 1A**) for which no resistance was detected on meropenem-containing plates (**Figure 2A,3A**) and was slightly underpredicted. The population fits did not describe the very late (from 215h), small regrowth (∼2 log_10_CFU/mL) for PAΔmexR challenged with 2g Q8h, which was not associated with resistance emergence at 239h and might be explained by biological variability (**Figure 1D,2D,3D**). For the 6g/day CI against PAΔDΔmexR, the model well described total bacterial counts on CAMHA for two biological replicates with a slight overprediction for the third replicate for which regrowth occurred slightly later (**Figure 1E**).

**Table 4:** Population parameter estimates for meropenem against isogenic strains of *P. aeruginosa*.

| Parameter | Symbol (unit) | Estimate (relative SE %) |
| --- | --- | --- |
| **Maximum population size** | $\log_{10}\text{CFU}_{\max}$ | 10.40 (0.43)<sup>c,f,g</sup> |
| | ($\log_{10}\text{CFU/mL}$) | 10.68 (0.34)<sup>a,b,d,e</sup> |
| **Mean generation time** | MGT (min) | 39.83 (1.01) |
| **Maximum killing effect** | $\text{KMAX}_{\text{MEM}} (\text{h}^{-1})$ | 1.67 (1.81) |
| **MEM concentration needed to achieve 50% of maximum killing** |  |  |
| Resistant | $\text{KC}_{\text{MEM}_R} (\text{mg/L})$ | 3.98 (2.70) |
| Intermediate | $\text{KC}_{\text{MEM}_I} (\text{mg/L})$ | 1.64 (7.34) |
| Susceptible | $\text{KC}_{\text{MEM}_S} (\text{mg/L})$ | 0.02 (4.42) |
| **Hill coefficient** | Hill | 0.56 (4.28) |
| **The effect of the *oprD* mutation on the rate of meropenem influx** | FOD1 | 0.21 (5.68) |
| **The maximum effect of the *mexR* mutation on the rate of meropenem efflux** | Vmax | 32.31 (2.14) |
| **MEM concentration at which efflux is at 50% of Vmax** | Km (mg/L) | 7.65 (2.47) |
| **The effect of the *ampD* mutation on the inactivation of meropenem** | FAD | 2.11 (4.00) |
| **$\log_{10}$ fraction of bacteria growing on meropenem-containing CAMHA<sup>h</sup>** |  |  |
| **3x MIC** |  |  |
| Resistant | $\log_{10} \text{FR}_{R3}$ | -3.68 (1.69)<sup>a</sup>, -0.45 (8.59)<sup>b</sup>,<br>-0.30 (15.12)<sup>c,d,e</sup>, -0.43 (7.66)<sup>f,g</sup> |
| Intermediate | $\log_{10} \text{FR}_{I3}$ | -5.14 (1.28)<sup>a</sup>, -3.86 (1.29)<sup>b</sup>,<br>-4.31 (1.90)<sup>c,d,e</sup>, -3.00 (1.66)<sup>f,g</sup> |
| Susceptible | $\log_{10} \text{FR}_{S3}$ | -7.60 (0.96)<sup>a</sup>, -7.05 (0.40)<sup>b</sup>,<br>-6.62 (0.44)<sup>c,d,e</sup>, -8.68 (0.32)<sup>f,g</sup> |
| **5x MIC** |  |  |
| Resistant | $\log_{10} \text{FR}_{R5}$ | -4.12 (1.60)<sup>a</sup>, -0.79 (9.27)<sup>b</sup>,<br>-0.58 (5.02)<sup>c,d,e</sup>, -1.37 (5.70)<sup>f,g</sup> |
| Intermediate | $\log_{10} \text{FR}_{I5}$ | -5.63 (1.00)<sup>a</sup>, -4.50 (0.70)<sup>b</sup>,<br>-5.30 (1.47)<sup>c,d,e</sup>, -4.54 (1.28)<sup>f,g</sup> |
| Susceptible | $\log_{10} \text{FR}_{S5}$ | -7.77 (0.46)<sup>a</sup>, -8.31 (0.54)<sup>b</sup>,<br>-7.47 (0.42)<sup>c,d,e</sup>, -9.43 (0.68)<sup>f,g</sup> |
| **Standard deviation of additive residual error on a $\log_{10}$ scale** |  |  |
| Total population | $\text{SD}_{\text{cfu}}$<br>($\log_{10}\text{CFU/mL}$) | 0.49 (3.65) |
| Population on 3x MIC meropenem CAMHA | $\text{SD}_{\text{cfu}3}$<br>($\log_{10}\text{CFU/mL}$) | 0.56 (5.88) |
| Population on 5x MIC meropenem CAMHA | $\text{SD}_{\text{cfu}5}$<br>($\log_{10}\text{CFU/mL}$) | 0.68 (6.09) |

<sup>a</sup>PAO1, <sup>b</sup>PAΔAD, <sup>c</sup>PAOD1, <sup>d</sup>PAΔmexR, <sup>e</sup>PAΔDΔmexR, <sup>f</sup>PAOD1ΔmexR, <sup>g</sup>PAOD1ΔD, <sup>h</sup>only one estimate was required per meropenem agar plate concentration across strains.

**Table 5:**
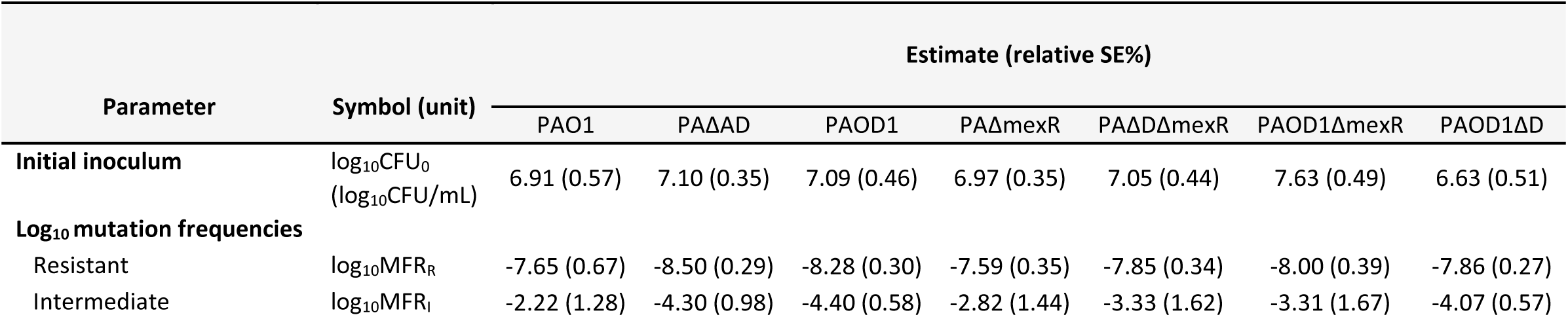
Population parameter estimates per strain for meropenem against isogenic strains of *P. aeruginosa*.

## Discussion

For carbapenems including meropenem, traditional PK/PD targets of 40% *f*T_>MIC_ from preclinical studies, and 70% *f*T_>MIC_ from clinical studies, have been linked to maximum bacterial killing and clinical cure, respectively (43). Targets of 100% *f*T_>MIC_ and 100% *f*T_>4-6×MIC_ have been proposed for severe infections in critically ill patients (44), and suppressing resistance emergence, respectively (45, 46). Across our HFIM studies, %*f*T_>MIC_ alone could not predict bacterial outcomes. Notably, attaining 98% *f*T_>MIC_ (2g Q8h) suppressed regrowth and resistance emergence of PAΔmexR, whereas even 100% *f*T_>5×MIC_ (6g/day CI) could not achieve this outcome against PAOD1 and PAΔDΔmexR, despite all three strains having the same meropenem MIC. Furthermore, attaining 100% *f*T_>MIC_ (6g/day CI) achieved extensive killing of PAOD1ΔD, but much less killing of PAOD1ΔmexR, despite the same MIC. Thus, we demonstrated that the PK/PD relationship was more complex than %*f*T_>MIC_. Even at the same MIC, the baseline resistance mechanism might affect the meropenem regimen required to suppress regrowth and resistance emergence.

Our results are also instructive regarding the effect of two pre-existing resistance mechanisms working together to influence bacterial response, including resistance emergence during meropenem exposure. The lowest CI and Q8h regimens of meropenem achieved substantial bacterial killing and suppressed resistance emergence against both PAO1 and PAΔAD. Thus, the *ampD* mutation that causes an increase in AmpC β-lactamase concentrations had little effect on bacterial outcome as a single resistance mechanism in PAΔAD compared to PAO1. This is not surprising since the *ampD* mutation has not been shown to substantially impact carbapenem effectiveness (16). Additionally, WGS revealed *oprD* mutational inactivation and MexAB-oprM overexpression through mutation of its negative regulators *mexR* or *nalD* were the primary mechanisms of meropenem resistance amplified during therapy. In contrast, mutations in *ampD* (or other *ampC* regulators such as *dacB* or *ampR*) were not amplified by meropenem exposure, indicating AmpC overexpression plays a minor role in constraining meropenem effectiveness. On the other hand, frequently selected mutations in aminoacyl tRNA synthetases likely reflect fitness cost compensatory mutations of the inactivation of *oprD*, coding for a porin involved in basic amino acids uptake (47). The bacterial response changed substantially when the *ampD* mutation was paired with a mutation resulting in increased efflux pumps (*mexR*) in PAΔDΔmexR as compared to PAΔmexR. Similarly, addition of the *ampD* mutation to the oprD mutant enhanced resistance mechanisms. While 12g/day CI suppressed resistance emergence of PAOD1 over 10 days, this regimen resulted in substantial resistance emergence in PAOD1ΔD from ∼3 days onwards. This highlights the importance of the interplay between multiple resistance mechanisms.

In contrast to %*f*T_>MIC_, our QSP model well characterized bacterial responses across all strains simultaneously, by accounting not only for the effects of the different baseline resistance mechanisms but also their interplay. Among the single mutants, the *oprD* mutation had the largest effect on meropenem resistance, followed by *mexR*. Similar to our study findings, OprD porin inactivation was reported to be the resistance mechanism conferring the highest level of meropenem resistance in *P. aeruginosa* clinical isolates (48, 49). Moreover, the model well described the effects of combinations of resistance mechanisms in double mutant strains, which notably included the observed enhancement effects of *ampD* plus *mexR* mutations. Current mechanism-based (or semi-mechanistic) models commonly include patients’ PK characteristics; however most do not directly include bacterial biological characteristics such as resistance mechanisms present. Typically, available models fit data from one, or up to three, strains in the same dataset. If multiple strains are present in the model, the differences in response to studied antibiotic(s) are frequently characterized by different parameter estimates for drug effect for each strain (21). These drug effect parameters are most often not informed by bacterial characteristics; therefore, are not predictable. In contrast, our novel model was informed by the underlying knowledge of key bacterial characteristics and allowed for quantification of the effects of pre-existing resistance mechanisms on antibiotic success and characterization of the interaction between resistance mechanisms, with the same parameter estimates of drug effect across strains.

Our results suggest the resistance mechanisms of the isogenic strains (*oprD*, *mexR*, *ampD*) when combined with a second mutation reduced bacterial killing, thus allowing for selection and amplification of pre-existing mutants with additional resistance mutations, which were responsible for driving regrowth and resistance emergence. The constant concentrations of such subpopulations in the control arms of each strain (detectable from 1-2 days onwards when the total population had reached a plateau) strongly suggests they were present prior to treatment. Furthermore, there was overlap between the mutations seen in the colonies from the control arm and the colonies from the treated arms for most strains, consistent with the assumption of amplification of pre-existing resistant mutants. The *aspS* mutation in the PAOD1ΔmexR control colony can be considered equivalent to the *lysS* mutations in the treatment arms (50, 51). In PAΔmexR a commonly occurring large genomic deletion in *galU* caused resistance in a control mutant but was not selected by treatment because of its higher fitness cost compared to the *oprD* mutation which was selected by treatment (52). Thus, the sequencing results (in addition to the baseline MFs) supported the model assumption that pre-existing resistant subpopulations drove the regrowth. In the model, the baseline resistance mutations decreased meropenem concentrations at the target site in the periplasmic space, which allowed for selection and amplification of pre-existing mutants with increased resistance to meropenem.

Standard, culture-dependent antimicrobial susceptibility testing is labour-intensive and often requires several days, which may delay selection of appropriate therapy (53, 54). Thus, many patients are still being treated empirically, which often leads to poor outcomes and increased resistance (55, 56). Currently, with the increasing burden of antimicrobial resistance, rapid WGS-based diagnostic testing is an option that could allow for identification of the infecting pathogen including detection of resistance genes (57). By identifying resistance characteristics, a therapy adapted to the individual pathogen (and patient) is an exciting prospect. Coupling our novel model with genomic analysis could represent a step towards optimizing antibiotic regimens against *P. aeruginosa* infections (58).

It is important to consider strengths and limitations of the study. To the best of our knowledge, it is the first to investigate response to meropenem in the context of *P. aeruginosa* resistance mechanisms in a HFIM, followed by development of a novel QSP model. The study included isogenic strains with a range of different resistance mechanisms and varying susceptibility to meropenem, and examined the impact of mode of administration of meropenem (CI *versus* intermittent infusion) on bacterial outcomes and resistance emergence. The tested isogenic strains reflected important resistance mechanisms common in clinical isolates, and were challenged with clinically relevant meropenem regimens over 10 days (6, 59). Additionally, inclusion of biological replicates ensured reproducibility of results. The quantification of resistant subpopulations on 3× and 5×MIC meropenem-containing CAMHA allowed for investigation of resistance emergence and their incorporation in the model. Genome sequencing of resistant colonies provided detailed insights into mechanisms of resistance emergence during meropenem exposure. In our study, meropenem in the periplasmic space was not measured. However, utilizing isogenic strains allowed for accurate estimation of the effects of different resistance mechanisms on bacterial response to meropenem for each strain and regimen. Since the effects are driven by the meropenem concentrations at the site of action (periplasmic space) in the isogenic strains compared to PAO1, an accurate estimate of the permeation rate or the nominal values of concentrations at the site of action were not required for the purpose of the current analysis. There is a paucity of information available on the rate of outer membrane permeability of meropenem in *P. aeruginosa* (60); as more becomes available the model could be extended. Similar to other *in vitro* models, the HFIM lacks an immune system; therefore, the observed bacterial responses may best reflect the meropenem effects as could be seen in immunosuppressed patients. Lastly, *P. aeruginosa* can also attain carbapenem resistance through acquisition of carbapenemase genes (encoding meropenem-deactivating enzymes) by horizontal gene transfer or mobile elements such as plasmids (61). Isogenic strains with increased carbapenemase production were not included in the present study which focused on chromosomal resistance mechanisms; however, future studies could address that topic.

In summary, we examined the effect of pre-existing *P. aeruginosa* resistance mechanisms on the effectiveness of simulated meropenem exposure profiles, reflecting PK of critically ill patients, in the HFIM, and developed a QSP model. The pharmacodynamic response was more complex than could be explained by %*f*T_>MIC_. We showed that, even for strains with the same MIC, the baseline resistance mechanisms affected bacterial outcomes and the meropenem regimens required to suppress regrowth and resistance emergence. The model well characterized the responses of seven isogenic strains simultaneously. The novel QSP model represents a step towards better understanding of the complicated relationship between antibiotic, bacterial characteristics and resistance emergence during treatment. This investigation provided a potential future approach towards enabling optimized antibiotic therapy. Future studies could explore other resistance mechanisms (including acquired resistance) and antibiotics in monotherapy and combinations.

## Sequence information

European Nucleotide Archive project number PRJEB71060.

