## Supplemental materials for "Elucidating effects of single and multiple resistance mechanisms on bacterial response to meropenem by quantitative and systems pharmacology modeling and population genomics"

### Supplementary methods

*Baseline mutation frequencies*

The baseline $\log_{10}$ MF reflected the proportion of pre-existing resistant mutant subpopulations present in the total population prior to antibiotic treatment. It was calculated as the difference between the $\log_{10}$ colony forming units (CFU)/mL on antibiotic-containing agar plates and the $\log_{10}$ CFU/mL on antibiotic-free agar. Briefly, for each isogenic strain, a bacterial suspension was grown overnight in sterile CAMHB at 37°C. Samples of the bacterial suspension with the desired inoculum target of $10^8$CFU/mL were plated on antibiotic-free (100µL) and 3× and 5×MIC meropenem-containing (200µL) CAMHA plates. Antibiotic-free plates were incubated for 24h, whereas antibiotic-containing plates were incubated for 48h at 37°C, followed by manual counting.

*Bioanalytical methods for pharmacokinetic validation*

Meropenem concentrations were determined by a validated liquid chromatography-tandem mass spectrometry (LC−MS/MS) assay, using a Shimadzu Nexera UHPLC-system coupled with a triple quadrupole 8030 Shimadzu mass spectrometer detector. For preparation of calibration standards and spiked quality control (QC) samples, appropriate amounts of meropenem standard solutions were added to CAMHB. A 50-µL aliquot of the supernatant from the samples was deproteinized by 70µL of 0.5%v/v trichloroacetic acid containing the internal standard (cefepime). After thorough mixing, the samples were centrifuged for 5min at 15,000g, and the supernatant was analysed by liquid chromatography-tandem mass spectrometry (LC−MS/MS).

For determination of the meropenem concentrations, 5µL of each PK sample was chromatographed on a Phenomenex Synergi column (150 × 2mm, 4µm, 80 Å; Phenomenex, Torrence, CA, U.S.A.) and eluted by using the isocratic condition with mobile phases A: water with 0.1%v/v formic acid and B: acetonitrile with 0.1%v/v formic acid, 81:19 (A:B) for 4min. Flow was 0.4mL/min and column oven temperature was 40°C. An electro-spray ionization source interface operating in positive-ion mode was used for the mass spectrometric multiple reaction monitoring. Quantitation of meropenem was performed using selected reaction monitoring of the transition of *m/z* 384.0 to 144.2 and 114.4 (reference ion) for meropenem, m/z 481.0 to 395.9 for cefepime. Under these conditions, meropenem and the internal standard were eluted after ~2min, and the total analysis time for one sample was 4min. LabSolutions software (version 5.6) was used for evaluation of the chromatograms. No interference was observed for the study drug or the internal standard. Five standard concentrations in the range of 0.2–40mg/L and 3 QC concentrations (0.5, 2, 30mg/L) were included. All data on intraday and interday precision and accuracy are presented in the Table below. The lower limit of quantification (LLOQ) was 0.2mg/L.

**Table:** Intraday and interday precision and accuracy of the LC-MS/MS assay

| **Parameter** | **Meropenem concentration (mg/L)** | | | |
|---|---|---|---|---|
| | **0.2 (LLOQ)** | **0.5 (QC-L)** | **2 (QC-M)** | **30 (QC-H)** |
| Precision (% CV) | | | | |
| Intraday (n = 3) | 2.22 | 4.00 | 4.54 | 9.47 |
| Interday (n = 12) | 5.48 | 3.69 | 3.90 | 7.47 |
| Accuracy (% bias) | | | | |
| Intraday (n = 3) | -5.22 | -6.22 | -0.54 | 0.81 |
| Interday (n = 12) | -2.25 | -6.02 | -1.10 | -0.66 |

#### *Genome sequencing and analysis of meropenem resistance mechanisms*

Total genomic DNA was extracted with a commercial kit (High Pure PCR Template Preparation Kit (Roche Diagnostics). Indexed paired-end libraries were then prepared with the Illumina DNA Prep Library Preparation Kit and sequenced on an Illumina NovaSeq 6000 sequencer (Illumina Inc, USA) with an SP flow cell, resulting in 300bp paired-end reads.

In order to study the emerging resistance mechanisms to meropenem, a variant calling analysis was performed following previously defined and validated protocols with slight modifications (1). Briefly, reads were mapped to the *P. aeruginosa* PAO1 reference genome (GenBank Accession Number NC_002516.2) with Bowtie2 v2.2.6 and pileup and raw files were obtained by using SAMtools v0.1.16 and PicardTools v1.140, using the Genome Analysis Toolkit (GATK) v.3.4-46 for realignment around InDels. For colonies, single nucleotide polymorphisms (SNPs) were extracted from the raw files if they met the following criteria: a quality score (Phred-scaled probability of the samples reads being homozygous reference) of at least 50, a root-mean-square (RMS) mapping quality of at least 25 and a coverage depth of at least 3 reads, excluding all ambiguous variants; and MicroInDels were extracted from the totalpileup files when meeting the following criteria: a quality score of at least 500, a RMS mapping quality of at least 25 and support from at least one-fifth of the covering reads. These filtered files were converted to vcf and then annotated with SnpEff v4.2. Totalpileup files obtained from total population samples were also converted to vcf and annotated with SnpEff v4.2, and the percentage of reads supporting the encountered variant (No. reads supporting the variant / No. total reads in that position) was extracted from the totalpileup files. Gene absence was also investigated in all saved colonies using Bowtie2 v2.2.6 and the SeqMonk software (https://www.bioinformatics.babraham.ac.uk/projects/seqmonk/). Nucleotide changes already present in the parental strain PAO1 and/or its isogenic derivatives (PAΔAD, PAOD1, PAΔmexR, PAΔDΔmexR, PAOD1ΔmexR and PAOD1ΔD) were disregarded.

Genomic sequences have been deposited in the European Nucleotide Archive under project number PRJEB71060.

*Quantitative and Systems Pharmacology (QSP) modeling*

The growth and replication of *P. aeruginosa* strains were described by a life-cycle growth model where bacteria are present in two states; state 1 – preparing for replication and state 2 – immediately before replication (2, 3). The final model included three pre-existing (present prior to antibiotic treatment) bacterial subpopulations with different susceptibilities to meropenem. The bacterial killing effect of meropenem on each subpopulation (susceptible, intermediate and resistant) was described as a direct effect model with a Hill function (4-6).

The viable counts on drug-free and meropenem-containing plates across all strains were modeled simultaneously, using the Monte Carlo Importance Sampling Algorithm (pmethod = 4) in parallelized S-ADAPT facilitated by SADAPT-TRAN (7, 8). Biological replicates were recorded separately in the modeling dataset. The S-ADAPT objective function (-1×log-likelihood), agreement between observed data and population predicted profiles, standard diagnostic plots, coefficients of correlation and biological plausibility of the parameter estimates were considered in assessing candidate models. Observations of <10 colonies per agar plate (<2$\log_{10}$ CFU/mL) were fitted *via* a previously developed residual error model (3). Otherwise, an additive error model was used to fit the viable counts for both antibiotic-free and meropenem-containing agar plates (3).

The model was developed with the assumption that the meropenem concentration at the site of action, *i.e.* intracellular periplasmic space, ($C_{intra}$) was dependent on the presence of different resistance mechanisms in the bacteria. The differential equation for $C_{intra}$ was defined as:

$$\frac{d(C_{intra})}{dt} = k_{eq} \cdot Fk_{in} \cdot C_{hf} - k_{eq} \cdot Fk_{out} \cdot C_{intra} \qquad (1)$$

The equilibration rate constant $k_{eq}$ was assumed to be fast (1000 $h^{-1}$), which meant that the equilibration between meropenem in the broth and the periplasmic space was almost instantaneous. The permeability rate of carbapenems for *P. aeruginosa* has been reported to be fast; however, while published data are available for *Klebsiella pneumoniae* and *Enterobacter cloacae*, there is still a lack of data for *P. aeruginosa*, as stated by Kim *et al.* (9, 10). Importantly, the model was not sensitive to this choice, as smaller values for $k_{eq}$ (*e.g.* 20 $h^{-1}$) did not affect parameter estimates or model performance. The $C_{hf}$ was the extracellular concentration of meropenem in broth in the HFIM experiment. The resistance mechanisms affected the concentration ratio between $C_{intra}$ and $C_{hf}$ at steady-state. The terms $Fk_{in}$ and $Fk_{out}$ reflected the fraction coefficients of the equilibration rate constant which characterized the rate of influx of meropenem from the extracellular to the periplasmic space ($Fk_{in}$) and the loss of meropenem from the periplasmic space ($Fk_{out}$), essentially determining the net flux of meropenem at equilibrium. Each of the three resistance mechanisms had a distinctive $Fk_{in}$ (porin loss) or $Fk_{out}$ (efflux pump overexpression, AmpC hyperproduction) estimated by the model and presented in the Table below.

**Table:** Distinctive $Fk_{in}$ and $Fk_{out}$ for each strain**.**

| **Strain** | **$Fk_{in}$** | **$Fk_{out}$** |
|---|---|---|
| PAO1 | 1 | 1 |
| PAΔAD | 1 | FAD |
| PAOD1 | FOD1 | 1 |
| PAΔmexR | 1 | FmexR |
| PAΔDΔmexR | 1 | FAD × FmexR |
| PAOD1mexR | FOD1 | FmexR |
| PAOD1ΔD | FOD1 | FAD |

**FOD1:** The effect of the ***oprD*** mutation on the rate of meropenem influx
**FmexR:** The effect of the ***mexR*** mutation on the rate of meropenem efflux
**FAD:** The effect of the ***ampD*** mutation on the inactivation of meropenem

In the model, PAO1 was assumed to have normal equilibration of meropenem and was treated as baseline, where $Fk_{in}$ and $Fk_{out}$ were set to 1 (*i.e.* 100%), while strains with *mexR* and/or *ampD* mutations were assumed to have increased removal of meropenem, decreasing $C_{intra}$. Strains with *oprD* mutation were modeled as having reduced entry of meropenem, also decreasing $C_{intra}$. The model assumed MexAB-OprM efflux pumps to be saturable and follow Michaelis–Menten kinetics, as suggested previously (11-13). The effect of the *mexR* mutation on the rate of meropenem efflux (FmexR) had a limit of 1 (baseline PAO1 effect) and was equal to:

$$FmexR = \frac{V_{max}}{K_m + C_{intra}} \tag{2}$$

In equation ( 2 ), $V_{max}$ represented the maximum value of FmexR and $K_m$ the $C_{intra}$ at which the system achieved 50% of FmexR.

The distinctive fraction coefficient of the equilibration rate constant, which characterized the rate of meropenem loss from the periplasmic space ($Fk_{out}$) for strain PAΔDΔmexR, was defined as a multiplication of the two effects that the resistance mechanisms present (resulting from the *ampD* and *mexR* mutations) had on the rate of meropenem removal. During model development, addition of the effects of these two resistance mechanisms was considered as an alternative. However, the latter model had substantially lower predictive performance and did not adequately characterize the time-course of bacterial outcomes for all strains with *ampD* and *mexR* mutations; therefore, the former model structure was selected.

*Bacterial killing by meropenem*

The bacterial killing effect of meropenem on the susceptible (predominant) bacterial subpopulation was described as:

$$KILL_{MEM\_S} = \frac{KMAX_{MEM} \cdot Cintra^{HILL}}{Cintra^{HILL} + KC_{MEM\_S}{}^{HILL}} \qquad (3)$$

where $KMAX_{MEM}$ was the maximum direct killing effect (a single maximum killing rate constant for all strains), $KC_{MEM}$ the $C_{intra}$ causing 50% of the maximum killing effect and HILL was the Hill coefficient, which described the steepness of the concentration-effect curve. The effect of meropenem on intermediate and resistant bacterial subpopulations were described similarly. Different estimates of $KC_{MEM}$ for the three different subpopulations (S – susceptible, I - intermittent and R – resistant) were required, but were the same for all strains.

The differential equations for the concentrations of the susceptible bacterial subpopulation in states 1 and 2 were:

$$\frac{d(CFU)_{S1}}{dt} = REP \cdot k_{21} \cdot CFU_{S2} - k_{12S} \cdot CFU_{S1} - KILL_{MEM_S} \cdot CFU_{S1} \qquad (4)$$

$$\frac{d(CFU)_{S2}}{dt} = -k_{21} \cdot CFU_{S2} + k_{12S} \cdot CFU_{S1} - KILL_{MEM_S} \cdot CFU_{S2} \qquad (5)$$

In the above equations, $k_{12}$ was the first-order growth rate constant describing the transition of bacteria from state 1 to state 2, and was equivalent to ln2/(MGT/60) with MGT being the mean generation time (min). The $k_{21}$ was the first-order replication rate constant (assumed to be very fast, not rate limiting) and was fixed to 50 $h^{-1}$ as described previously (2). The intermittent and resistant bacterial subpopulations were modeled similarly.

The REP function ensured that $CFU_{all}$ did not exceed the maximum population size ($CFU_{max}$) (14).

$$REP = 2 \cdot (1 - \frac{CFU_{all}}{CFU_{all}+CFU_{max}}) \quad (6)$$

At a low $CFU_{all}$, REP approached 2, representing a 100% probability of successful replication. As $CFU_{all}$ approached the maximum population size ($CFU_{max}$), REP approached 1, which represents a 50% probability of successful replication with the total viable count remaining constant and bacteria continuing to transition between states 1 and 2.

*Resistant bacterial populations on antibiotic-containing agar*

The viable counts on meropenem-containing agar (3x and 5x meropenem MIC) were co-modeled with the viable counts on drug-free agar. The fractions of different bacterial subpopulations, that were able to grow on meropenem-containing agar, were estimated as described previously (15), however with only one parameter estimate needed per meropenem concentration in agar across strains (instead of separate estimates for each strain). The fractions of susceptible ($FR_S$), intermediate ($Fr_I$) and resistant ($FR_R$) bacterial subpopulations were included. The viable counts of bacteria on meropenem-containing agar were described as:

$$CFU_{on\_3\times MIC} = FRS_{S3} \cdot CFU_S + FR_{I3} \cdot CFU_I + FR_{R3} \cdot CFU_R \quad (7)$$

$$CFU_{on\_5\times MIC} = FR_{S5} \cdot CFU_S + FR_{I5} \cdot CFU_I + FR_{R5} \cdot CFU_R \quad (8)$$

where $CFU_S$, $CFU_I$ and $CFU_R$ were the concentrations of bacterial subpopulations (susceptible, intermediate and resistant respectively) in both states (state 1 and 2, as described above).

### SUPPLEMENTARY FIGURES

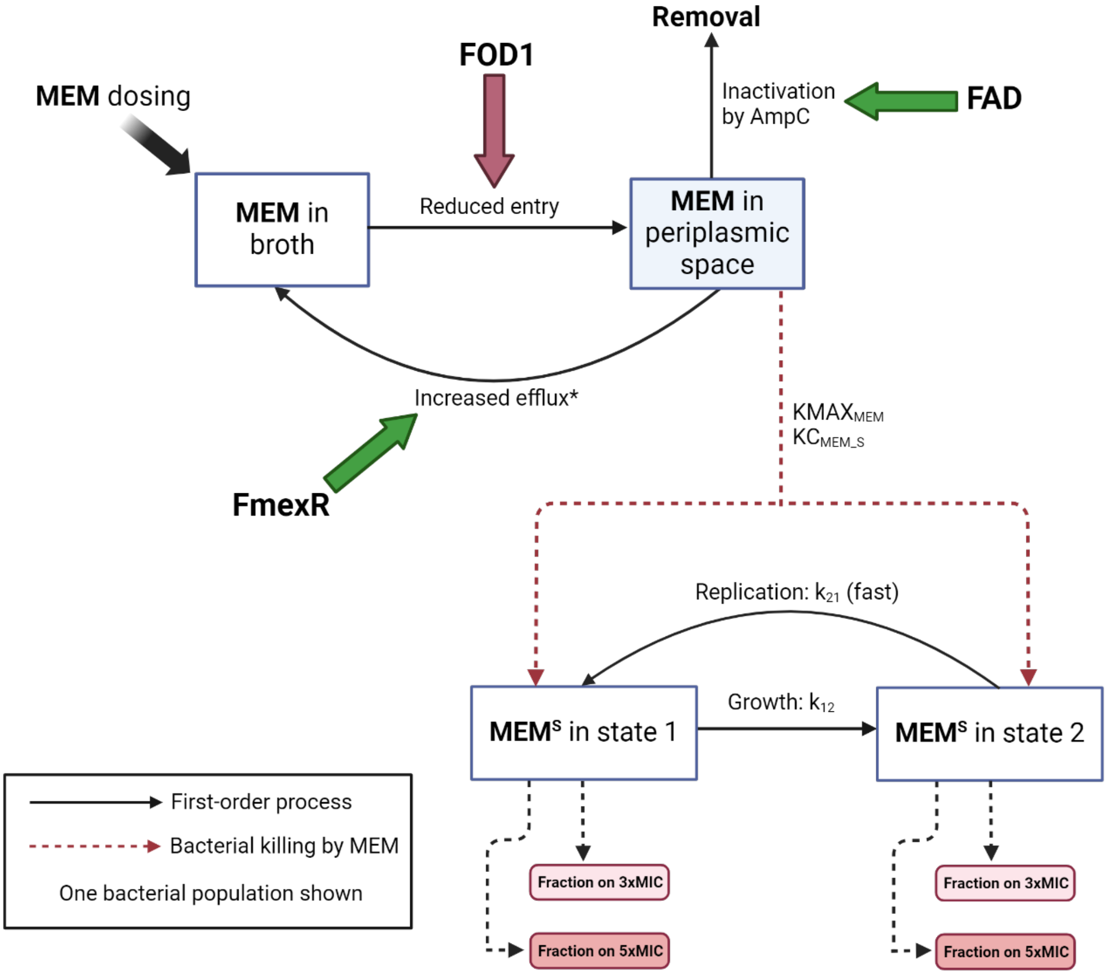


**Figure S1:** A schematic representation of the meropenem (MEM) MBM for $MEM^S$ (bacterial population susceptible to MEM). MEM intermediate and resistant populations, which are not shown, have substantially higher fractions of their bacterial populations growing on 3xMIC and 5xMIC agar. FOD1: the effect of the *oprD* mutation on the rate of MEM influx; FmexR: the effect of the *mexR* mutation on the rate of MEM efflux; FAD: the effect of the *ampD* mutation on the inactivation of MEM. The maximum killing rate constant $KMAX_{MEM}$ and associated $KC_{MEM\_S}$ causing 50% of $KMAX_{MEM}$ are explained in Table 5. *The amount of MEM that effluxes back to the broth was considered negligible and would not affect the MEM concentrations in broth; therefore, was not included in the model.

### Supplementary results

**Table S1:** The average baseline $Log_{10}$ mutant frequencies (MF) of seven isogenic strains of *P. aeruginosa* against meropenem.

| Strain | Concentration of meropenem in CAMHA (mg/L) | $Log_{10}$ MF |
|---|---|---|
| PAO1 | 3 (3×MIC) | -8.20 |
| | 5 (5×MIC) | < -8.89 |
| PAΔAD | 6 (3×MIC) | -6.94 |
| | 10 (5×MIC) | < -8.97 |
| PAΔmexR | 12 (3×MIC) | -8.12 |
| | 20 (5×MIC) | < -8.87 |
| PAOD1 | 12 (3×MIC) | -6.95 |
| | 20 (5×MIC) | < -8.94 |
| PAΔDΔmexR | 12 (3×MIC) | -5.54 |
| | 20 (5×MIC) | -6.43 |
| PAOD1ΔmexR | 48 (3×MIC) | -6.65 |
| | 80 (5×MIC) | -8.83 |
| PAOD1ΔD | 48 (3×MIC) | -9.13 |
| | 80 (5×MIC) | < -8.74 |

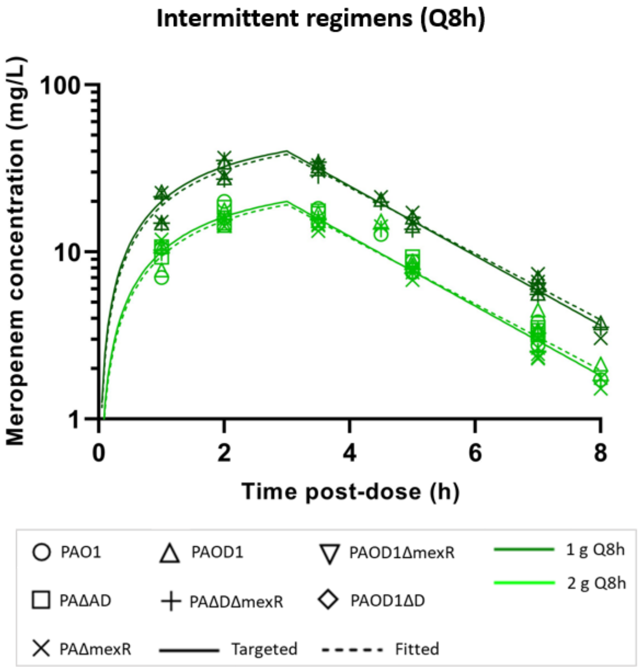


**Figure S2:** Targeted (solid lines) and observed (symbols) meropenem concentrations in the HFIM. The observed concentrations and fitted profiles (broken lines) reflected the targeted PK profiles of meropenem in critically ill patients with normal renal function.

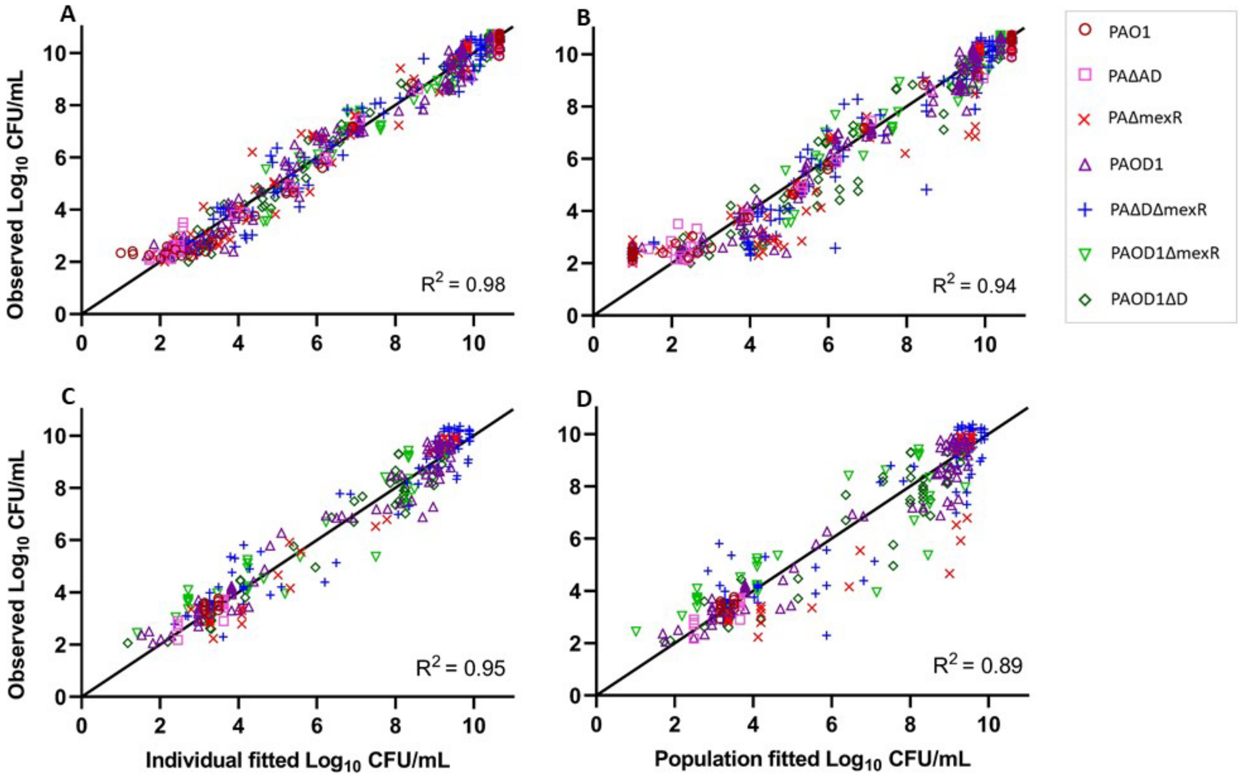


**Figure S3:** Observed vs individual- and population-fitted counts (A and B on drug-free CAMHA, and C and D on meropenem containing CAMHA) of *P. aeruginosa* isogenic strains for the developed MBM.
